# PKR and the Integrated Stress Response drive immunopathology caused by ADAR1 mutation

**DOI:** 10.1101/2020.11.30.405498

**Authors:** Megan Maurano, Jessica M. Snyder, Caitlin Connelly, Jorge Henao-Mejia, Carmela Sidrauski, Daniel B. Stetson

## Abstract

Mutations in *ADAR*, the gene that encodes the ADAR1 RNA deaminase, cause numerous human diseases, including Aicardi-Goutières Syndrome (AGS). ADAR1 is an essential negative regulator of the RNA sensor MDA5, and loss of ADAR1 function triggers inappropriate activation of MDA5 by self-RNAs. However, the mechanisms of MDA5-dependent disease pathogenesis *in vivo* remain unknown. Here, we introduce a knockin mouse that models the most common *ADAR* AGS mutation in humans. These *Adar*-mutant mice develop lethal disease that requires MDA5, the RIG-I-like receptor LGP2, type I interferons, and the eIF2α kinase PKR. We show that a small molecule inhibitor of the integrated stress response (ISR) that acts downstream of eIF2α phosphorylation prevents immunopathology and rescues the mice from mortality. These findings place PKR and the ISR as central components of immunopathology *in vivo* and identify new therapeutic targets for treatment of human diseases associated with the ADAR1-MDA5 axis.

## Introduction

The RIG-I-like receptors (RLRs) MDA5 and RIG-I are important for the rapid detection of viral RNAs and the initiation of the type I interferon (IFN) response, which restricts viral replication and spread (Goubau et al., 2013). However, this mode of defense comes with the risk of potential recognition of self-RNAs and chronic activation of the IFN response. Negative regulation of these nucleic acid sensors is therefore essential to prevent inappropriate recognition of endogenous RNAs by the RLRs (Crowl et al., 2017).

The RNA-editing enzyme Adenosine Deaminase Acting on RNA 1 (ADAR1) edits endogenous double-stranded RNAs (dsRNAs) to prevent chronic activation of MDA5 and IFN production in response to self-RNA (George et al., 2016; Liddicoat et al., 2015; Mannion et al., 2014; Pestal et al., 2015). ADAR1 is expressed as two distinct isoforms: the constitutively expressed 110 kDa ADAR1 p110, and the IFN-inducible 150 kDa ADAR1 p150 (Patterson and Samuel, 1995). ADAR1 edits mRNA coding sequences (Hartner et al., 2004), micro RNAs (Yang et al., 2006), viral RNAs (Tenoever et al., 2007), and, most frequently, inverted repeats of SINE retroelements in noncoding RNA regions (Osenberg et al., 2010). The inosine products of ADAR1 deamination are read as guanosine by the translational machinery, so RNA editing by ADAR1 can result in non-synonymous coding changes. Inosines also disrupt the dsRNA structures necessary for MDA5 activation (Ahmad et al., 2018; Liddicoat et al., 2015). *In vitro*, MDA5 binds to these same regions - predominantly inverted repeats of SINEs - in cells lacking ADAR1 (Ahmad et al., 2018; Chung et al., 2018; Ishizuka et al., 2019). The p150 isoform of ADAR1 is specifically responsible for MDA5 regulation, whereas the p110 isoform contributes to multiorgan development independent of MDA5 regulation (Pestal et al., 2015).

Over 200 distinct mutations in the *ADAR* gene have been identified that cause numerous human diseases associated with a type I IFN response, including Aicardi-Goutières syndrome (AGS), Bilateral Striatal Necrosis (BSN), and Dyschromatosis Symmetrica Herediteria (DSH; Hayashi and Suzuki, 2013; Livingston et al., 2014; Rice et al., 2012). Despite considerable progress in understanding the molecular mechanisms of AGS and related diseases, these conditions remain untreatable and incurable, underscoring the need to identify new therapeutic targets for intervention. Three mouse models of ADAR1 mutation have been instrumental in defining the relationship between ADAR1 RNA editing and the MDA5-MAVS pathway: *Adar*-null mice that lack both ADAR1 isoforms (Hartner et al., 2004; Hartner et al., 2009; Wang et al., 2004), an *Adar p150* knockout mouse that lacks ADAR1 p150 but retains ADAR1 p110 (Ward et al., 2011), and a knockin *Adar* point mutation that disrupts deaminase activity but retains ADAR1 protein expression (Liddicoat et al., 2015). Moreover, CRISPR targeting of the *ADAR* gene in human cells has been used to further explore consequences of ADAR1 loss (Ahmad et al., 2018; Chung et al., 2018; Pestal et al., 2015), and ADAR1 blockade was recently identified as a potential strategy to enhance innate immune responses in tumor cells (Ishizuka et al., 2019). However, characterization of the ADAR1-MDA5 regulatory axis *in vivo* has been hampered by the embryonic lethality caused by the null alleles of *Adar* in mice, which can only be rescued by simultaneous disruption of the genes that encode MDA5 or MAVS (Liddicoat et al., 2015; Mannion et al., 2014; Pestal et al., 2015). Thus, study of the relationship between ADAR1 and MDA5 in live mice has been impossible using current models.

To address these limitations and to enable the identification of AGS disease mechanisms *in vivo*, we generated a knockin mouse that models the most common *ADAR* allele found in AGS: a nonsynonymous point mutation that converts a proline to an alanine at position 193 in the human ADAR1 protein (P193A; P195A in mice). This mutation is located within the Zα domain that is unique to the p150 isoform, and it is present at a remarkably high allele frequency of ~1/360 in humans of northern European ancestry (Crow et al., 2015; Rice et al., 2012). Using these mice, we show that *Adar P195A* paired with a null allele of *Adar* causes complete, postnatal, MDA5-dependent mortality. We further define essential requirements for the RLR LGP2, type I IFNs, and the eIF2α kinase PKR in disease progression. Finally, we show that therapeutic inhibition of the integrated stress response (ISR) downstream of PKR is sufficient to completely prevent disease. Together, these data reveal effector mechanisms downstream of MDA5 activation that contribute to immunopathology *in vivo*, with implications for treatment of human diseases caused by *ADAR* mutation.

## Results

### The *Adar P195A* knockin mouse model

Over 60% of AGS patients with *ADAR* mutations carry the P193A allele as a compound heterozygote with either a frameshift mutation or a mutation in the deaminase domain of ADAR1 (Crow et al., 2015; Rice et al., 2012). Interestingly, no AGS patients have been identified who are homozygous for *ADAR P193A*. We hypothesized that this was because homozygosity of P193A mutation would be incompatible with life, similar to the phenotype of total ADAR1 loss in mice (Hartner et al., 2004; Wang et al., 2004). To determine how this mutation impacts ADAR1 function and self-RNA detection, we used CRISPR targeting of fertilized mouse oocytes to generate mice carrying the orthologous P195A mutation in the endogenous *Adar* locus (Figure 1A). We intercrossed *Adar P195A/+* mice and identified live births of *Adar+/+, Adar P195A/+*, and *Adar P195A/P195A* mice at the expected Mendelian ratios (Figure 1B). We tracked the survival and weights of *Adar P195A/+* and *Adar P195A/P195A* mice and found them to be indistinguishable from wild type controls (Figure 1C, 1D). This suggests that the absence of known *ADAR P193A/P193A* AGS patients might be due to a lack of disease. We then measured expression of *Adar* mRNA in cerebellum and in bone marrow-derived macrophages (BMMs) and found that the *Adar P195A* mutation did not impact *Adar* mRNA levels in resting cells or after treatment with recombinant IFN beta (IFNβ; Supplemental Figure 1). Moreover, the protein expression levels, inducibility, and nuclear and cytosolic distribution of the ADAR1 p110 and p150 isoforms were unaffected by the *Adar P195A* mutation (Supplemental Figure 1).

**Figure 1.**
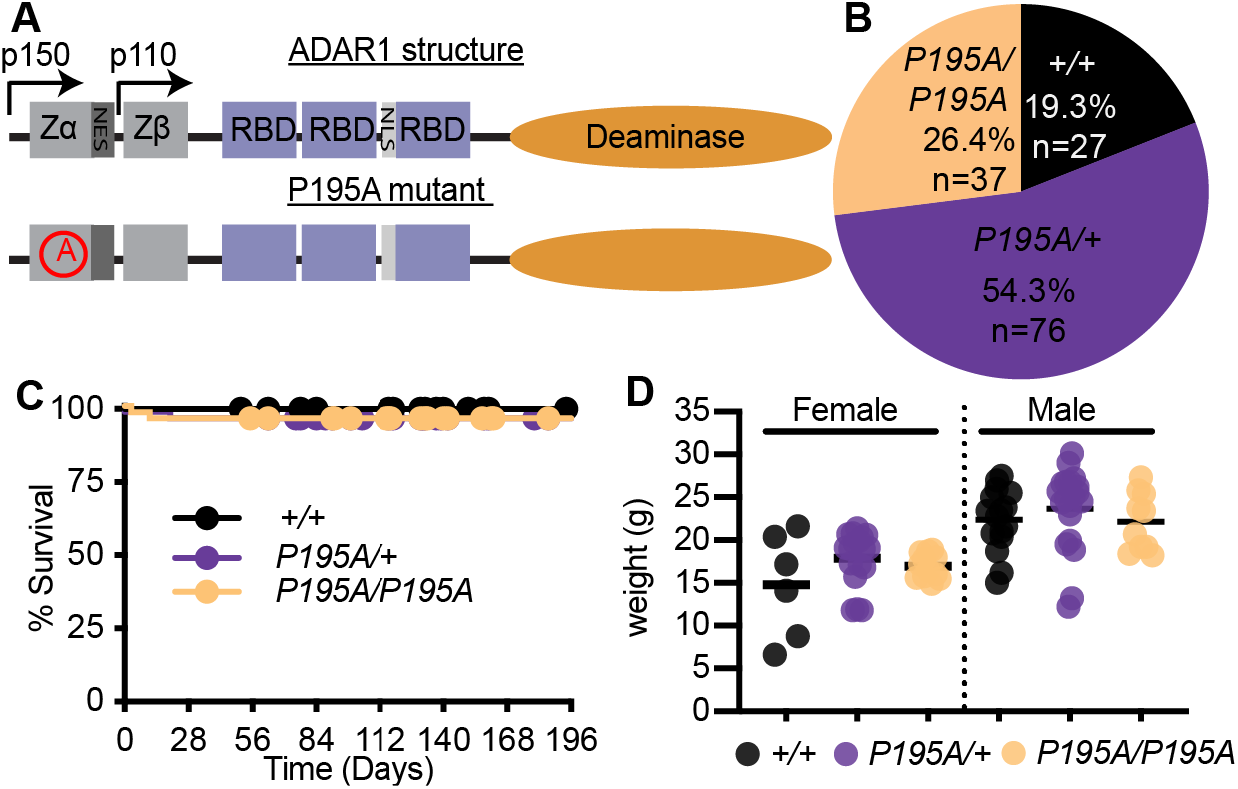
The *Adar P195A* knock-in mouse model. (A) Schematic of the structure of ADAR1, and the location of the P195A mutation. Zα, Zβ, = Z-DNA binding domains, NES = Nuclear export signal, NLS = nuclear localization signal. (B) Percentage of mice of the indicated genotype from intercrosses of *Adar P195A/+* mice. (C) Survival of *Adar +/+* (n=23), *Adar P195A/+* (n=47), *Adar P195A/P195A* (n=48) mice. (D) Weights of mice of the indicated *Adar* genotypes at 23 days of age. Bar represents the mean.

### Recapitulation of AGS patient genotypes causes severe disease in *Adar P195A* mice

Because the *ADAR P193A* mutation in AGS is invariably found as a compound heterozygote with a more severe *ADAR* allele (Crow et al., 2015; Rice et al., 2012), we intercrossed *Adar P195A/P195A* mice with either *Adar+/-* or *Adar p150+/-* mice to model the combinations of *ADAR* alleles found in AGS patients. We found that both *Adar P195A/-* mice and *Adar P195A/p150*- mice were born at frequencies that matched the recovery of heterozygous mice from crosses of the parental *Adar*-null and *Adar p150*-null alleles (Supplemental Figure 2A, 2B). However, and in stark contrast to the *Adar P195A/P195A* mice (Figure 1C), we noted complete postnatal mortality when the P195A mutation was paired with either the full *Adar* null allele or the *Adar p150*-null allele (Figure 2A, 2B). Mortality in the *Adar P195A/-* mice (median survival 21 days) progressed more rapidly than in the *Adar P195A/p150*- mice (median survival 40 days; Figure 2A, 2B). Next, we performed these same intercrosses of the *Adar P195A* mutation with the *Adar*-null and *Adar p150*-null alleles on an *Ifih1-/-* (MDA5 KO) background and observed complete rescue from mortality of both *Adar P195A/-Ifih1-/-* and *Adar P195A/p150-Ifih1-/-* mice (Figure 2C, 2D). Consistent with the essential contribution of MDA5 to disease, we found that the *Adar P195A/-* and *Adar P195A/p150*- mice were severely runted at weaning compared to littermate controls, and that this runting was entirely MDA5-dependent (Figure 2E, 2F). We also noted that heterozygosity for *Ifih1* delayed mortality, revealing a gene dosage-specific effect of MDA5 expression on disease (Supplemental Figure 2C, 2D). Taken together, these data demonstrate that the *Adar P195A* mice recapitulate the human *ADAR* genotypes found in AGS and develop severe disease that is driven by MDA5. More broadly, they represent the first *Adar*-mutant mice that are born with intact MDA5 signaling, allowing the dissection of disease mechanisms *in vivo*.

**Figure 2.**
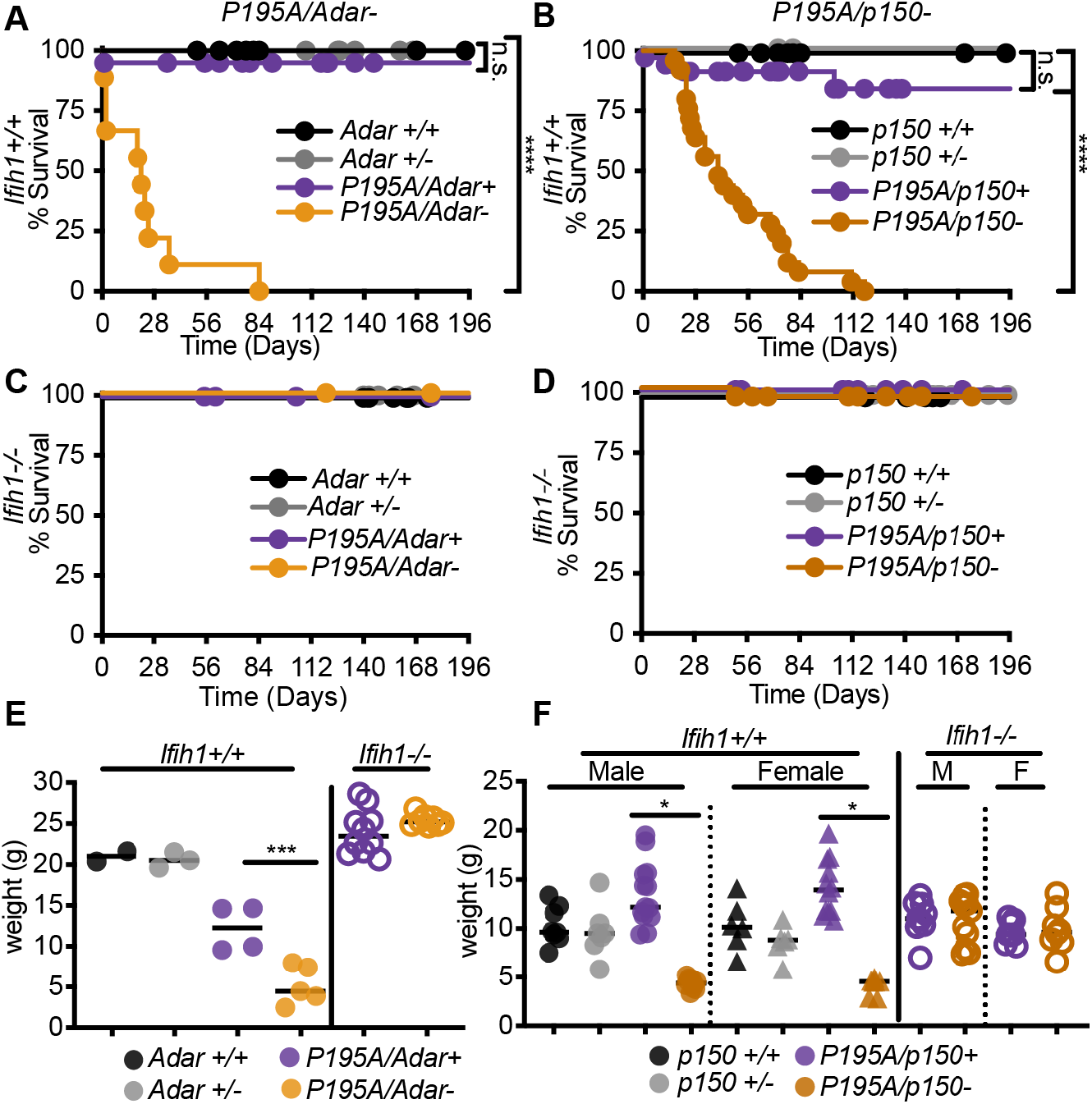
Recapitulation of AGS patient genotypes causes severe disease in *Adar P195A* mice. (A-B) Survival of *Ifihi1+/+* mice of the indicated genotypes: *Adar+/+* (n=14), *Adar+/-* (n=21), *Adar P195A/Adar+* (n=41), *Adar P195A/Adar*- (n=9); *Adar p150+/+* (n=15), *Adar p150+/-* (n=6), *Adar P195A/p150+* (n=34), *Adar P195A/p150*- (n=25). (C-D) Survival of *Ifihi1-/-* mice of the indicated genotypes: *Adar+/+* (n=18), *Adar+/-* (n=22), *Adar P195A/Adar+* (n=14), *Adar P195A/Adar*- (n=19); *Adar p150+/+* (n=17), *Adar p150+/-* (n=24), *Adar P195A/p150+* (n=25), *Adar P195A/p150*- (n=28). (E-F). Weights of mice of the indicated genotypes, measured at 23 days. Bars represent mean. Male and female mice are pooled in (E) because there was no significant difference by sex.

To better understand the causes of runting and mortality in this model, we performed necropsies on sex-matched *Adar P195A/p150*- and *Adar P195A/p150+* littermates to evaluate the pathologies associated with disease. We focused these and all additional analyses on the *Adar P195A/p150*- mice because they survived longer than *Adar P195A/-* mice (Figure 2A, 2B). In our initial assessment, the clearest histological defects were found in the kidney and liver. The kidney exhibited glomerular mesangial matrix expansion, and the liver exhibited extensive microvesicular cytoplasmic vacuolation that increased with age (Figure 3A–3C). Strikingly, and in contrast to the *Trex1-/-* mouse model of AGS that is driven by the cGAS-STING DNA sensing pathway rather than by MDA5-MAVS (Gall et al., 2012; Gao et al., 2015; Gray et al., 2015; Stetson et al., 2008), we found no evidence of inflammatory immune cell infiltrates in these tissues (Figure 3A–3C). We additionally identified dysregulated architecture of the spleens in *Adar P195A/p150*- mice (Figure 3C). We developed a histological scoring approach to quantitate these pathologies across several mice per genotype. *Adar P195A/p150*- mice had significantly higher pathological scores in all three organs, and these scores were normalized to control levels in *Adar P195A/p150-Ifih1-/-* mice (Figure 3D–3F). As an independent measure of liver function, we found that serum alkaline phosphatase (ALP) levels were elevated, and serum albumin levels were reduced in *Adar P195A/p150*- mice compared to controls (Figure 3G). We next grew primary BMMs from *Adar P195A/p150*- mice on *Ifih1+/+* and *Ifih1-/-* backgrounds. The *Adar P195A/p150*-BMMs did not spontaneously express elevated levels of *Ifnb* mRNA, but treatment of these cells with recombinant IFNβ instigated robust, MDA5-dependent *Ifnb* transcription (Figure 3H), as has been previously observed in *ADAR*-null human cell lines (Ahmad et al., 2018). Together, these data demonstrate severe, MDA5-dependent pathologies and MDA5-mediated aberrant type I IFN expression in *Adar P195A/p150*- mice.

**Figure 3.**
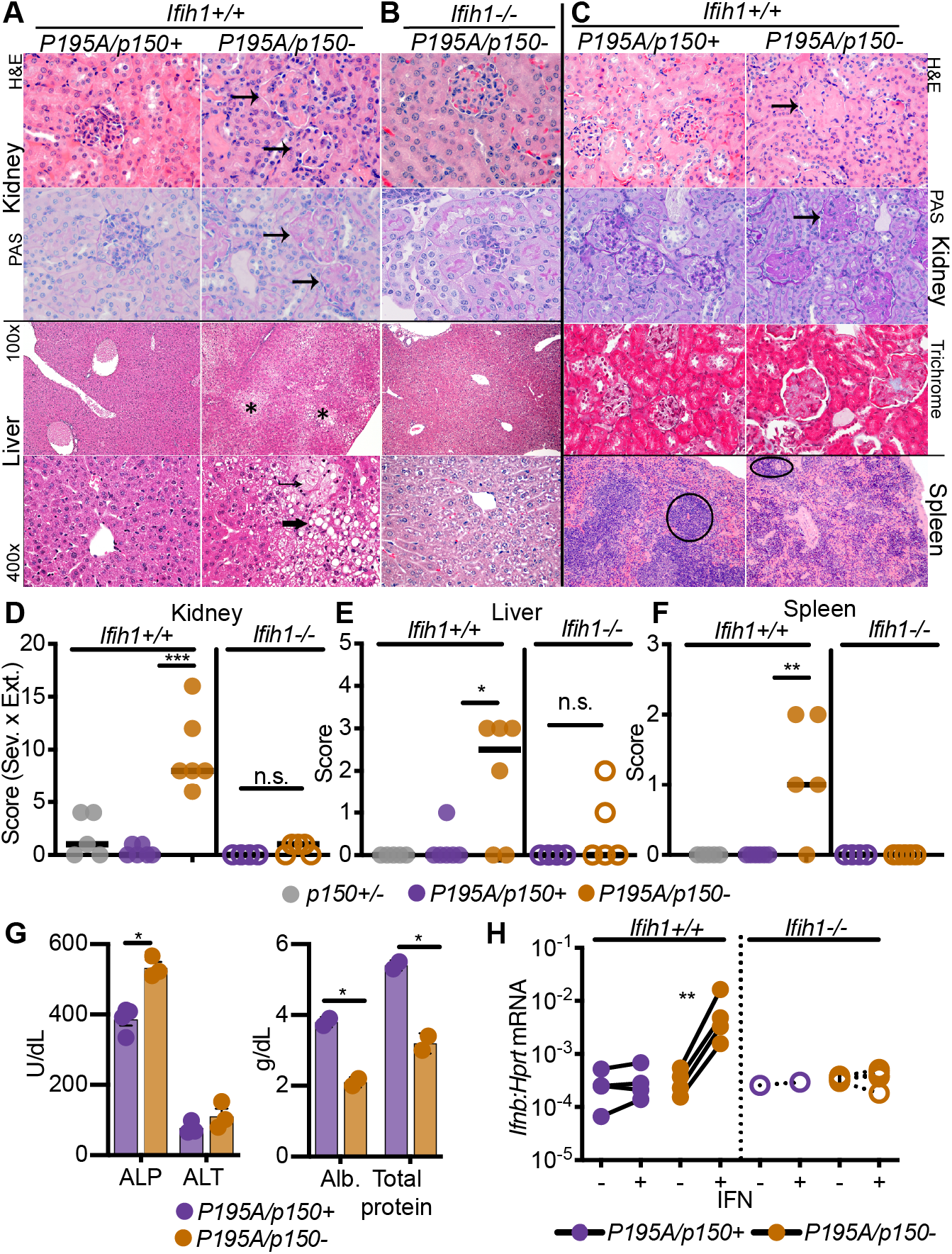
Organ-specific pathology in *Adar P195A/p150*- mice. (A) Representative histology of kidney and liver from *Adar P195A/p150*- mice measured at 23 days of age. (B) Representative histology of kidney and liver from *Adar P195A/p150-Ifih1-/-* mice measured at 23 days of age. (C) Representative histology of kidney, liver, and spleen from *Adar P195A/p150*- mice measured at 147 days of age. (D-F) Histological scores in the kidney, liver, and spleen, measured at 23 days of age. (G) ALP, ALT, Albumin, and protein measured in the serum of 23 day old mice. Bars represent mean and SEM. (H) *Ifnb* transcript measured by qRT-PCR in BMMs of the indicated genotypes, with and without 24 hours of treatment with recombinant mouse IFNβ.

We performed mRNA-Seq analyses comparing age-matched *Adar P195A/p150*- mice and *Adar P195A/p150+* controls to quantitate global changes in gene expression caused by the *Adar P195A* mutation. We analyzed liver and kidney because of the specific pathologies we uncovered in these tissues (Figure 3), and we additionally included cerebellum to compare changes in gene expression in the brains of these mice. Focusing on well-curated interferon-stimulated genes (ISGs), specifically genes in the GO term ‘response to type I interferon’, we carried out a gene set enrichment analysis and found significant up-regulation of this gene set in all three tissues (Figure 4A–4C). The extent of the ISG signatures varied among tissues, with cerebellum showing the most significant increases in ISG expression (adjusted p=0.007), followed by kidney (adjusted p=0.003) and then liver (adjusted p=0.01; Figure 4A–4C). We then performed quantitative RT-PCR of selected ISGs in all three of these tissues, comparing additional *Adar P195A/p150*- mice and *Adar P195A/p150+* controls, on both *Ifih1+/+* and *Ifih1-/-* backgrounds. Consistent with the complete rescue from mortality and pathology in *Adar P195A/p150-Ifih1-/-* mice (Figures 2 and 3), we found that the significantly elevated ISG expression was also entirely MDA5-dependent (Figure 4D–4F). Thus, the *Adar P195A/p150*- mice recapitulate the MDA5-dependent ISG signatures found in the blood and cells of AGS patients (Uggenti et al., 2019).

**Figure 4.**
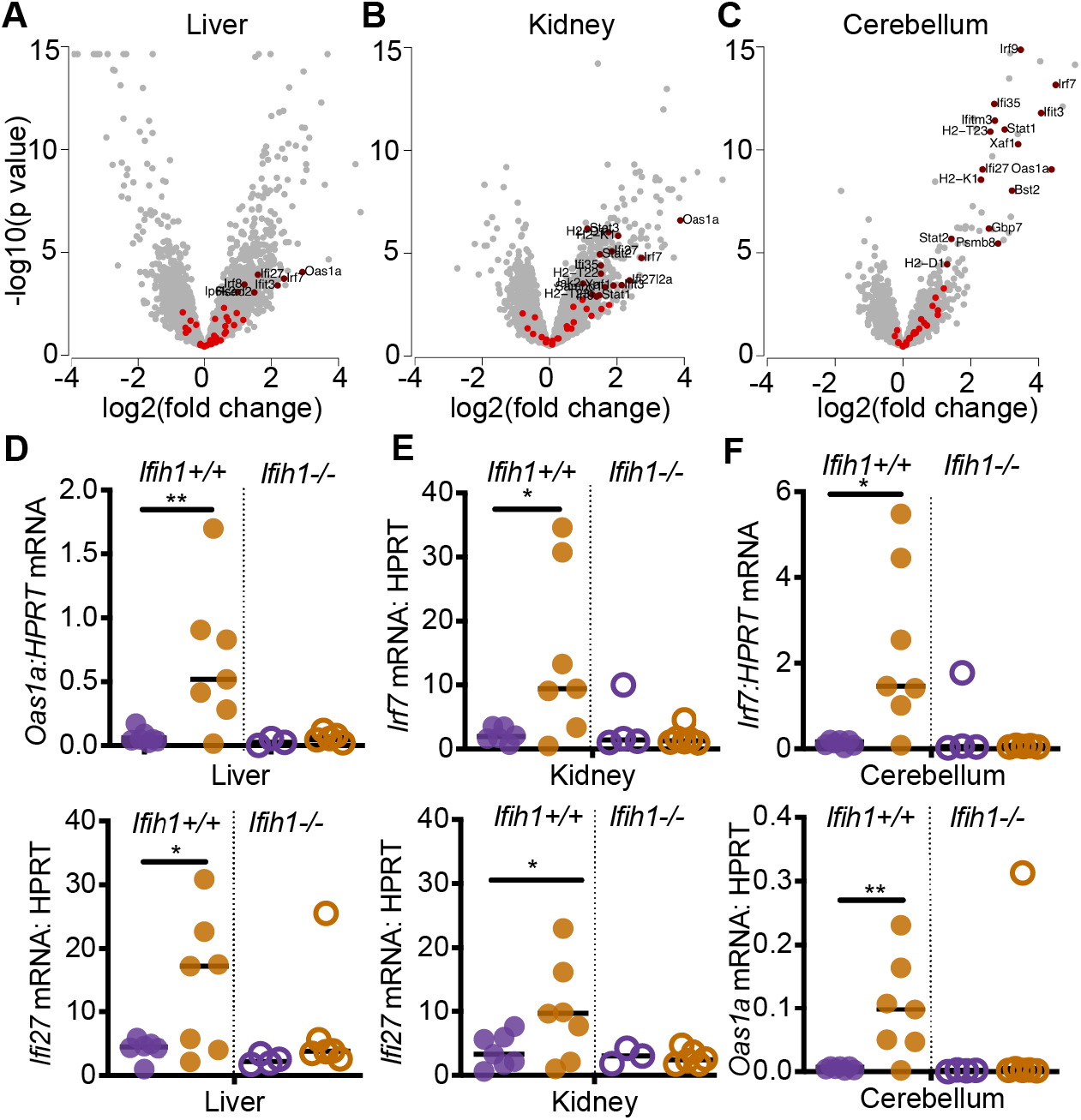
MDA5-dependent interferon signature in *Adar P195A/p150*- mice. (A-C) Expression data for ISGs, defined by the GO term ‘response to type I interferon,’ were evaluated in the liver, kidney, and cerebellum of 23 day old *Adar P195A/p150*- mice, plotting the log2 fold change over matched *Adar P195A/p150+* control mice. ISGs that were not significantly changed are shown in bright red; significant expression changes are shown in dark red. (D-F) Expression of ISGs identified in A-C, measured by TaqMan qPCR, in 23 day old mice of the indicated genotypes.

### Genetic dissection of the *Adar P195A/p150*- phenotype *in vivo*

The development of postnatal, lethal, MDA5-dependent disease in *Adar P195A/p150*- mice allowed us to further characterize how aberrant MDA5 signaling results in pathology *in vivo*. To do this, we performed a series of crosses to test stringently the contributions of additional signaling pathways, monitoring survival, weights, and ISG signatures. We started these analyses by examining the contribution of the third RIG-I-like receptor, LGP2, which is encoded by the *Dhx58* gene. The role of LGP2 in antiviral immunity is less well understood than that of MDA5 and RIG-I, in part because LGP2 lacks the Caspase Activation and Recruitment domain (CARD) that is essential for RIG-I and MDA5 signaling through MAVS. Moreover, a role for LGP2 in any immune pathology has not been previously reported. However, numerous studies have demonstrated that LGP2 interacts with MDA5 (Deddouche et al., 2014), modulates the formation of MDA5 filaments on dsRNAs (Bruns et al., 2014; Duic et al., 2020), and is important for antiviral responses to RNA viruses that specifically activate MDA5 (Satoh et al., 2010). We intercrossed *Adar P195A/P195A* mice with *Adar p150+/-* mice on a *Dhx58-/-* background, analyzing *Adar P195A/p150*- mice and comparing them to *Adar P195A/p150+* littermate controls. We found that LGP2 deficiency completely rescued the *Adar P195A/p150*- mice from mortality, restored their weights to normal, and prevented the ISG signature in cerebellum, liver, and kidney (Figure 5A–5C). These findings place LGP2, together with MDA5, at the apex of the signaling pathway that links ADAR1 dysfunction to disease.

**Figure 5.**
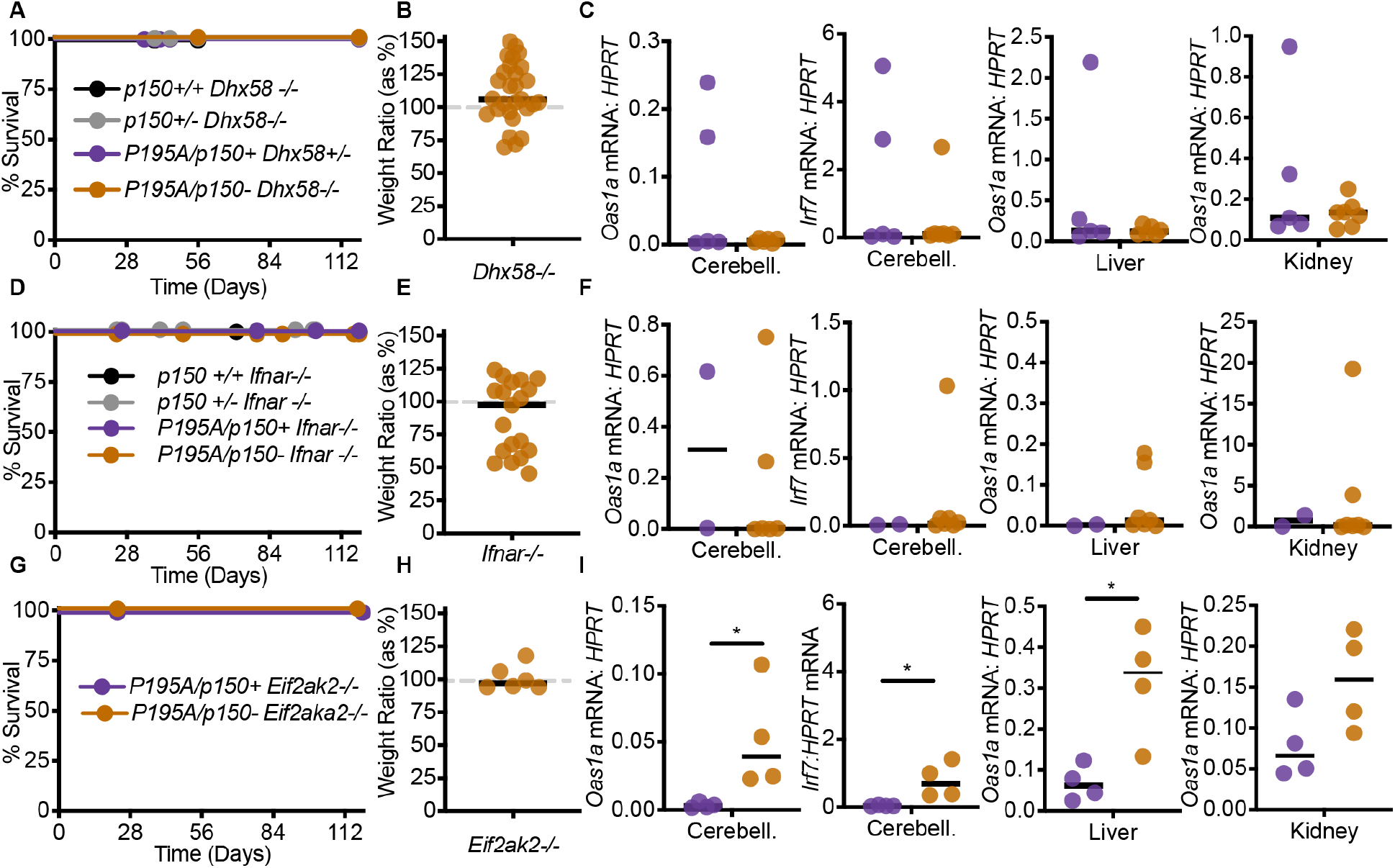
Genetic dissection of the *Adar P195A/p150*- phenotype *In vivo*. (A) Survival of *Dhx58-/-* mice of the indicated genotype: *Adar p150+/+* (n=3), Adar *p150+/-* (n=8), *Adar P195A/p150+* (n=5), *Adar P195A/p150*- (n=12). (B) Weights, measured at 23 days, of *Adar P195A/p150-Dhx58-/-* mice, as a percentage of the average weight of age- and sex-matched *Adar P195A/p150+Dhx58-/-* control mice. (C) Expression of the indicated ISGs measured by TaqMan qRT-PCR, normalized to HPRT, in the cerebellum, liver, or kidney, comparing *Adar P195A/p150-Dhx58-/-* mice to *Adar P195A/p150+Dhx58-/-* controls. (D) Survival of *Ifnar-/-* mice of the indicated genotype: *Adar p150+/+* (n=3), *Adar p150+/-* (n=7), *Adar P195A/p150+* (n=4), *Adar P195A/p150*- (n=16). (E) Weights, measured at 23 days, of *Adar P195A/p150-Ifnar1-/-* mice, as a percentage of the average weight of age- and sex-matched *Adar P195A/p150+Ifnar1-/-* control mice. (F) Expression of the indicated ISGs measured by TaqMan qRT-PCR, normalized to HPRT, in the cerebellum, liver, or kidney, comparing *Adar P195A/p150-Ifnar1-/-* mice to *Adar P195A/p150+Ifnar1-/-* controls. (G) Survival of *Eif2ak2-/-* mice of the indicated genotype: *Adar P195A/p150+* (n=4), or *Adar P195A/p150*- (n=7). (H) Weights, measured at 23 days, of *Adar P195A/p150-Eif2ak2-/-* mice, as a percentage of the average weight of age- and sex-matched *Adar P195A/p150+Eif2ak2-/-* control mice. (I) Expression of the indicated ISGs measured by TaqMan qRT-PCR, normalized to HPRT, in the cerebellum, liver, or kidney, comparing *Adar P195A/p150-Eif2ak2-/-* mice to *Adar P195A/p150+Eif2ak2-/-* controls.

Next, we tested the importance of the ISG signatures in *Adar P195A/p150*- mice by crossing them to *Ifnar1-/-* mice that lack the type I interferon signature. We found that IFNAR1 deficiency also completely rescued all aspects of disease (Figure 5D–5F), similar to the rescue of *Trex1-/-* mice on an *Ifnar1-/-* background (Stetson et al., 2008).

We evaluated the *in vivo* contribution of the dsRNA-activated eIF2α kinase PKR (encoded by the *Eif2ak2* gene) to disease in *Adar P195A/p150*- mice. PKR is activated by similar dsRNA structures as those that activate MDA5/LGP2, and the activation of PKR in ADAR1-deficient cells has been clearly demonstrated in both mouse and human cell lines (Chung et al., 2018; George et al., 2016; Ishizuka et al., 2019; Li et al., 2010). Additionally, prior studies have identified an important role for PKR upstream of the activation of type I IFNs by MDA5 (Pham et al., 2016; Schulz et al., 2010). However, a specific contribution of PKR to *in vivo* immunopathology has not been described. We found that *Adar P195A/p150-Eif2ak2-/-* mice were completely protected from mortality and weight loss (Figure 5G–5H). However, the ISG signature remained significantly elevated in cerebellum and liver from these mice (Figure 5I). These findings directly implicate PKR in the disease caused by ADAR1 dysfunction *in vivo*, and they place PKR as an essential downstream effector, rather than an activator, of the MDA5/LGP2-dependent IFN response in this model.

We examined the contribution of the endonuclease RNase L to disease in the *Adar P195A/p150*- mouse model. RNase L is activated by the 2’-5’ oligoadenylate (2-5A) products of the dsRNA-activated OAS enzymes (Kristiansen et al., 2011; Zhou et al., 1993), which are IFN-inducible nucleotidyltransferases that resemble cGAS in overall structure and catalytic mechanism (Civril et al., 2013). Once activated by 2-5A, RNase L cleaves cellular and viral RNAs, which limits viral replication and restricts mRNA translation (Hornung et al., 2014). RNase L has been previously implicated as a key effector that mediates cell death downstream of *ADAR* disruption in human cell lines (Daou et al., 2020; Li et al., 2017). We found that RNase L deficiency had no impact on the mortality or weight loss in *Adar P195A/p150*- mice (Supplemental Figure 3), suggesting that the contribution of RNase L to disease in this model might be more subtle or cell type specific.

Together, this genetic dissection of the *Adar P195A/p150*- mouse model reveals new and essential contributors to *in vivo* immunopathology and places them in a hierarchy that links MDA5-and LGP2-dependent IFN responses to PKR-dependent effector mechanisms that drive disease.

### The ISR is responsible for pathology and mortality caused by *Adar* mutation

Based on the striking and complete rescue of PKR-deficient *Adar P195A/p150*- mice, we explored the contribution of PKR-dependent effector mechanisms to disease in more detail. PKR is one of four metazoan eIF2α kinases that couple diverse perturbations in cellular homeostasis to a program called the integrated stress response (ISR), which restricts new protein synthesis and results in the transcriptional induction of genes that can either restore homeostasis or cause cell death, depending on the strength and duration of the insult (Costa-Mattioli and Walter, 2020; Harding et al., 2003). The eIF2 GTPase is responsible for the delivery of the initiator methionyl tRNA to the ribosome that initiates mRNA translation at the AUG start codon. GTP hydrolysis releases eIF2 from the ribosome-mRNA complex, after which the eIF2 must be recycled from its GDP-bound inactive form into its GTP-bound active form in order to initiate a new round of mRNA translation (Hinnebusch and Lorsch, 2012). Phosphorylation of the a subunit of eIF2 on serine 51 prevents this recycling of the eIF2 complex by the guanine nucleotide exchange factor (GEF) eIF2B, and results in a reduction of most canonical mRNA translation initiation (Krishnamoorthy et al., 2001). However, certain mRNAs that contain unusual arrangements of AUG start codons in their 5’ untranslated regions become selectively translated in the context of eIF2α phosphorylation (Sachs et al., 1997). These include the transcription factor ATF4, which induces the expression of ISR-activated genes (Harding et al., 2000).

The ISR gene expression program has been extensively defined in cell lines, and *in vivo* in mice that harbor hypomorphic mutations in *Eif2b5*, one of the genes that encodes the 5-subunit eIF2B complex (Wong et al., 2019). *EIF2B* gene mutations in humans cause a lethal leukoencephalopathy called Vanishing White Matter disease (VWM) that is driven by a chronic ISR (Leegwater et al., 2001; van der Knaap et al., 2002). Using a curated ISR gene expression signature defined in the *Eif2b5*-mutant mouse model of VWM (Wong et al., 2019), we examined our mRNA-Seq data and found significant up-regulation of the ISR gene set in *Adar P195A/p150*- mice (Figure 6A-C). The ISR gene set was significantly elevated in liver (adjusted p=0.02) and kidney (adjusted p=0.07) but was not significantly increased in cerebellum (adjusted p=0.57; Figure 6A–6C). Taken together with the ISG analyses (Figure 3), the liver and kidney exhibited elevation of both ISG and ISR gene sets, but the cerebellum exhibited only the ISG signature (Figure 6A–6C).

**Figure 6.**
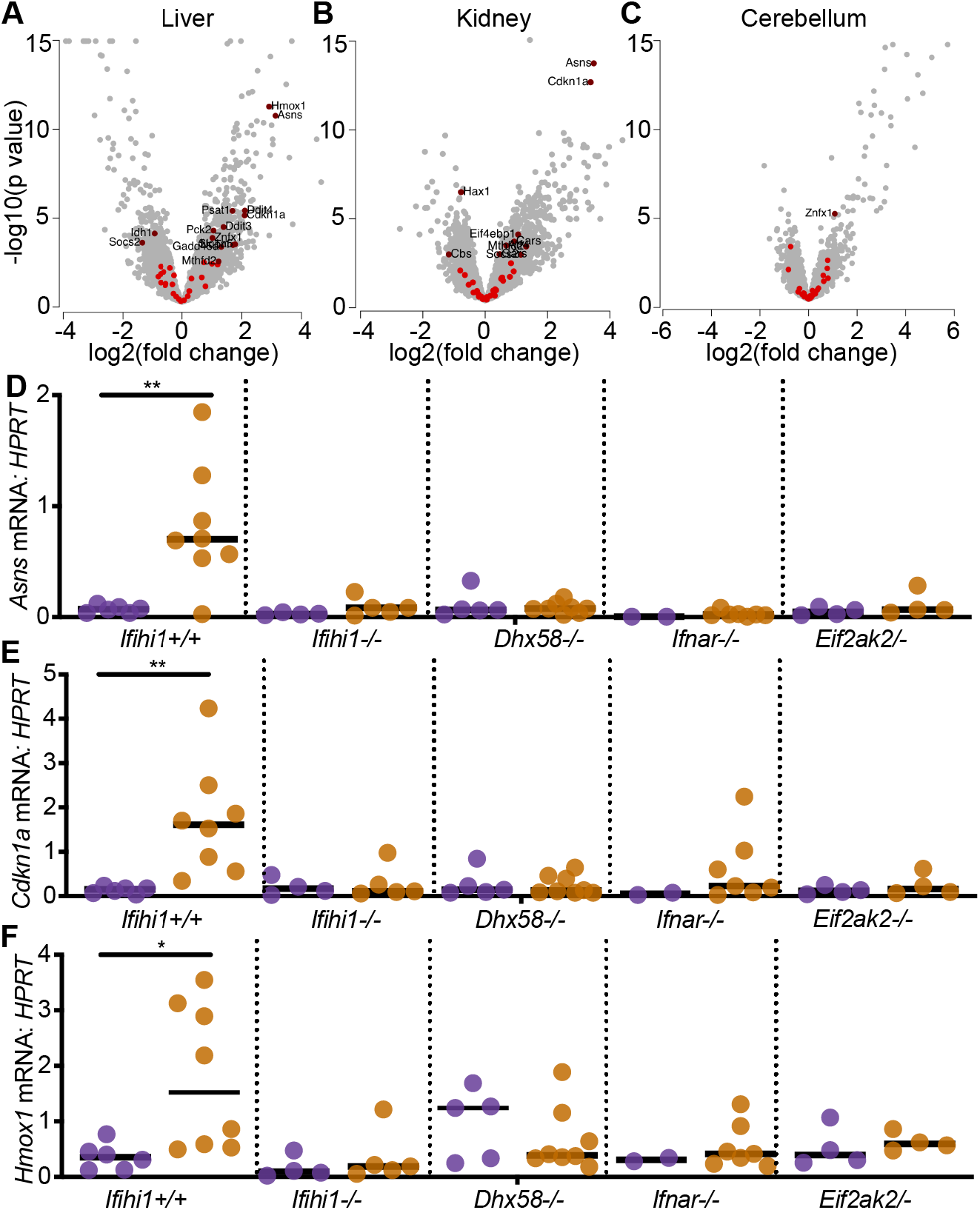
An ISR gene expression signature in *Adar P195A/p150*- mice. (A-C) Expression data for ISR gene set genes in the liver, kidney, and cerebellum of 23 day old *Adar P195A/p150*- mice, plotting the log2 fold change over control *Adar P195A/p150+* mice. ISR genes that are not significantly changed are shown in bright red; significant expression changes are shown in dark red. (D-F) Expression of ISR transcripts identified in the livers of rescued mice, measured by TaqMan qRT-PCR, and compared among the indicated genotypes. Each data point represents an individual mouse.

We selected three of the most robustly induced ISR genes in the livers of *Adar P195A/p150*- mice (*Asns, Cdkn1a*, and *Hmox1*), and evaluated their expression by quantitative RT-PCR, comparing the affected mutant mice to all of the rescued genotypes that we identified in Figures 2 and 5. We found that induction of ISR gene expression required MDA5, LGP2, IFNAR1, and PKR (Figure 6D–6F), which precisely mirrored the rescue from mortality and pathology in all of these genotypes, even more specifically than the ISG signature that remained elevated in PKR-deficient *Adar P195A/p150*- mice (Figure 5I).

A small molecule called Integrated Stress Response Inhibitor (ISRIB) stabilizes the eIF2B complex and activates its GEF function, rendering eIF2B less sensitive to the inhibitory effects of eIF2α phosphorylation (Sekine et al., 2015; Sidrauski et al., 2013; Sidrauski et al., 2015; Tsai et al., 2018; Zyryanova et al., 2018). More recently, a similarly potent and brain-penetrant analog of ISRIB with improved *in vivo* pharmacokinetics and pharmacodynamics called 2BAct was shown to prevent all aspects of VWM in a mouse model of *Eif2b5* mutation, including brain pathology and induction of ISR genes (Wong et al., 2019). We tested whether the ISR inhibitor 2BAct could impact disease in the *Adar P195A/p150*- mouse model. To do this, we formulated 2BAct into mouse chow (Wong et al., 2019), and we placed breeders on 2BAct-containing or control chow two days after timed matings. We maintained the mice on this regimen through birth and nursing of pups, and then continued treatment of the mice following weaning. We tracked survival and disease phenotypes for 125 days after birth, which is three times the median survival of unmanipulated *Adar P195A/p150*- mice (Figure 2). We found that dietary introduction of 2BAct nearly completely rescued the *Adar P195A/p150*- mice from mortality, and it restored the mice to normal weights (Figure 7A–7B). Moreover, 2BAct-treated *Adar P195A/p150*- mice were indistinguishable from controls when examined by tissue pathology in liver, kidney, and spleen, both at weaning and at the end of the 125-day treatment (Figure 7C–7D). Interestingly, the ISGs remained significantly elevated in 2BAct-treated *Adar P195A/p150*- mice (Figure 7E), but expression levels of the three ISR genes were restored to control levels (Figure 7F). Thus, we have shown that therapeutic amelioration of the ISR is sufficient to prevent mortality and pathology in this *Adar*-mutant mouse model, revealing an essential IFN-dependent effector mechanism that contributes to disease *in vivo*.

**Figure 7.**
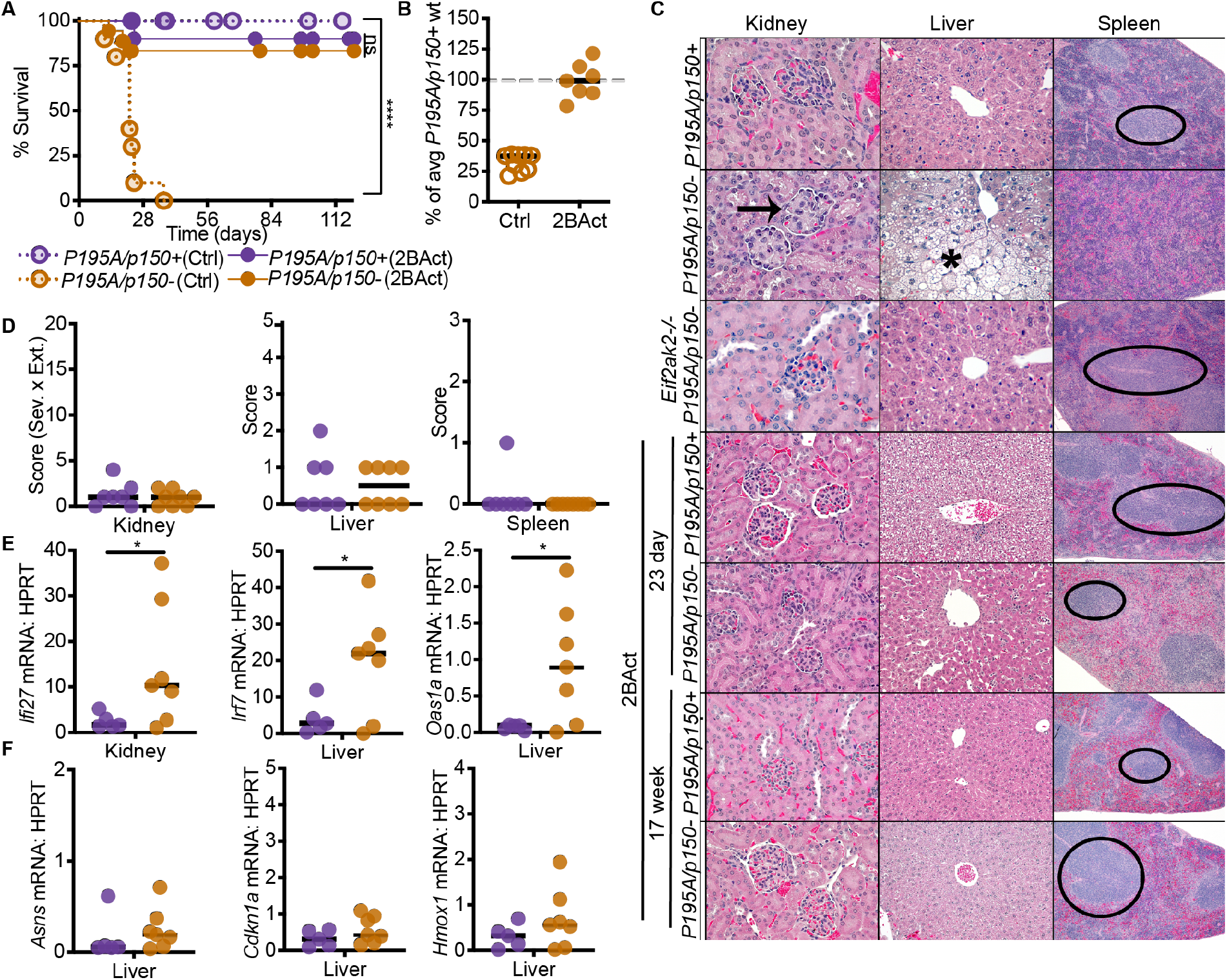
Pharmacological Inhibition of the ISR rescues *Adar P195A/p150*- mice. (A) Survival of mice on control chow: *Adar P195A/p150+* (n=23), *Adar P195A/p150*- (n=10); versus survival of mice on 2BAct chow: *Adar P195A/p150+* (n=17), *Adar P195A/p150*- (n=17). (B) Weights of *Adar P195A/p150*- mice on control chow or 2BAct chow, as a percentage of average weight of age- and sex-matched *Adar P195A/p150+* mice on control chow. (C) Representative histology of kidney, liver, and spleen of untreated, *Eif2ak2-/-*, and 2BAct-treated mice of the indicated genotypes. (D) Histological scores of the kidneys, livers, and spleens of 2BAct-treated mice at day 23. (E) ISG expression in the kidneys and livers of *Adar P195A/p150+* and *Adar P195A/p150*- mice treated with 2BAct. (F) ISR transcript expression in the liver of *Adar P195A/p150+* and *Adar P195A/p150*- mice treated with 2BAct. Bars represents the mean in all graphs for B-F.

## Discussion

We introduce a new mouse model of an AGS *Adar* mutation that develops postnatal, MDA5-dependent mortality. We use this model to delineate the genetic pathways responsible for pathology and we reveal a therapeutic approach that completely prevents disease, with implications for our understanding of disease mechanisms and targets for intervention in the human immune disorders caused by *ADAR* mutation.

Our studies of the *Adar P195A/p150*- mice offer new insights into the links between ADAR1 dysfunction, MDA5, and disease manifestations. First, we show that the RLR LGP2 is essential for the MDA5-mediated antiviral response in this model, revealing a new target for therapeutic intervention. Prior *in vitro* studies of the biochemical mechanisms of MDA5 filament formation on dsRNAs in the context of ADAR1 editing have focused exclusively on interactions between synthesized dsRNAs and purified recombinant MDA5 (Ahmad et al., 2018). Because LGP2 can modulate the size of MDA5 filaments and stabilize smaller MDA5-dsRNA complexes (Bruns et al., 2014; Duic et al., 2020), the size and composition of dsRNAs that are competent to trigger LGP2/MDA5 *in vivo* in the context of ADAR1 dysfunction might be distinct from those defined *in vitro*. Second, our findings reveal essential contributors to disease that were not appreciated in prior studies of the *Adar*-null alleles in mouse models. Specifically, neither IFNAR1 deficiency nor PKR deficiency rescued *Adar*-null mice to birth (Pestal et al, unpublished data; and Wang et al., 2004). However, we have now found that both IFNAR1 and PKR are essential for disease in the *Adar P195A/p150*- mouse model (Figure 5). This likely reflects the severity of the null alleles of *Adar* compared to the *Adar P195A* point mutation, together with the essential function of ADAR1 that is independent of its role in regulating MDA5 (Pestal et al., 2015). Unlike the *Adar*-null alleles, which eliminate both functions of ADAR1, the *Adar P195A* mouse model isolates the specific MDA5 regulatory roles of the p150 isoform of ADAR1.

By modeling the disposition of human *ADAR* AGS mutations in mice, we have established a genetic pathway that links ADAR1 dysfunction to disease. This pathway places MDA5 and now LGP2 as the initiating sensors that are required for detecting the self-RNAs that fail to be edited in *Adar P195A/p150*- mice. Next, the MDA5-and LGP2-dependent type I interferon response drives all aspects of disease pathology. Finally, we define PKR as an essential IFN-dependent effector that mediates the disease manifestations and mortality. Whereas PKR is clearly important for antiviral defense and it is targeted by virus-encoded antagonists (Elde et al., 2009), PKR has not previously been implicated in self RNA-mediated immune pathology *in vivo*.

We have found that the ISR inhibitor 2BAct, administered in food, prevents mortality and all aspects of disease pathology in *Adar P195A/p150*- mice. Moreover, we identify an ISR gene expression signature in affected mice that is more specific than the ISG signature in its correlation with genetic and therapeutic rescue. Specifically, the ISGs remain significantly elevated in PKR-deficient *Adar P195A/p150*- mice and in 2BAct-treated *Adar P195A/p150*- mice, but the loss of the ISR signature correlates perfectly with all of the genetic and therapeutic rescues that we have defined. These findings have a number of important implications. First, we reveal a new therapeutic path to treat human diseases caused by ADAR1 dysfunction that targets a specific effector mechanism while leaving other antiviral responses intact. Second, we directly implicate the ISR as the cause of immune pathology. Third, we identify a gene expression signature that could be harnessed to explore the contributions of the ADAR1-MDA5/LGP2-PKR axis to additional immune diseases in which IFNs and MDA5 have been implicated, including type I diabetes and systemic lupus erythematosus (Nejentsev et al., 2009; Robinson et al., 2011). More broadly, we speculate that the link between the ISR and immune pathology might also apply to conditions in which the ISR is activated by distinct mechanisms, including the endoplasmic reticulum stress sensor PERK, the heme sensor HRI, and the nutrient sensor GCN2 (Costa-Mattioli and Walter, 2020). All of these kinases share eIF2α as their principal target, and all induce the ISR. In other words, because the ISR contributes to inflammation and pathology downstream of PKR in this model of ADAR1 mutation, it might also drive inflammation downstream of diverse stresses that are not typically thought of as immune in origin. Indeed, ISR gene expression signatures have been identified in tissue samples from human patients with multiple sclerosis (MS; Way and Popko, 2016). Current models of MS envision the ISR as a cytoprotective mechanism that slows disease progression by preventing the loss of myelin-producing oligodendrocytes (Way and Popko, 2016). However, our findings raise the alternative possibility that the ISR might contribute to disease pathology. We propose that further study of protective versus pathogenic contributions of the ISR will illuminate new ways to classify and treat human immune diseases.

In summary, we have developed and characterized a mouse model of ADAR1 dysfunction that recapitulates human AGS genotypes, we have revealed new contributors to disease, and we have provided proof-of-concept for a therapeutic approach targeting the ISR as an essential contributor to immune pathology *in vivo*.

## Methods

### Mice

*Adar P195A* mutant mice were generated as previously described (Henao-Mejia et al., 2016), using the sgRNA target sequence GAAGGGGGAAACCTCCTTTGTGG, and the oligo DNA to replace the sequence, ATCAATCGTATTTTGTACTCCCTGGAAAAGAAGGGAAAGCTGCACAGAGGAAGGGGGAAACCTGCCTTGTGGAGCCTTGTGCCCTTGAGTCAGGCTTGGACTCAGCCCCCTGGAGTTGTGAATCCAGAT. In brief, C57BL6/J oocytes were microinjected with Cas9 complexed with gRNA for WT *Adar* sequence and ssDNA donor template containing the desired sequence. Oocytes were implanted into a pseudopregnant CD-1 females. Founder pups born were screened by the Surveyor assay. Mice positive by the Surveyor assay were genotyped using TaqMan SNP genotyping, primers: WT: AAACCT**CCT**/ZEN/TTGTGGAGCCTT, Mut: AAACCT**GCC**TTGTGGAGCCTT. Mice positive by SNP genotyping were then sequenced to confirm fidelity of the introduced mutation. Mice were bred to C57BL6/J mice to confirm germline transmission, and *Adar P195A/+* mice were then crossed to one another or to other genetically modified mice.

*Adar fl/fl* mice (Hartner et al., 2009) were generously provided by Stuart Orkin, and were bred to Cre-expressing mice to generate the *Adar*-null allele as described previously (Pestal et al., 2015). *Adar p150+/-* gametes (Ward et al., 2011) were generously provided by Michael Oldstone and made into mice as previously described (Pestal et al., 2015). *Ifih1-/-* mice (Gitlin et al., 2006) were generously provided by Michael Gale, Jr. *Dhx58-/-* mice (Suthar et al., 2012) were generously provided by Michael Gale, Jr. *Ifnar1-/-* mice (Muller et al., 1994) were backcrossed 14 generations to C57BL/6 (Kolumam et al., 2005). *Eif2ak2-/-* mice (Abraham et al., 1999), backcrossed at least 10 generations to C57BL/6J, were generously provided by Gökhan Hotamişligil. *Rnasel-/-* mice (Zhou et al., 1997) were generously provided by Robert Silverman. All mice were maintained in a specific pathogen-free (SPF) barrier facility at the University of Washington, and all experiments were done in accordance with the Institutional Animal Care and Use Committee guidelines of the University of Washington. Mouse weights were measured at 23 days of age.

### Histology and Pathology

For all histological analyses, matched littermate mice were euthanized in accordance with IACUC protocols by CO_2_ asphyxiation followed by cardiac puncture. Mice were washed with PBS and then fixed in 10% neutral-buffered formalin. Tissues were routinely processed, embedded in paraffin, cut into approximately 4 μm sections and hematoxylin and eosin stained. Slides of kidney, liver, and spleen were also stained with periodic acid-Schiff. Additionally, kidney sections were stained with Congo Red and Masson’s trichrome.

Tissues were evaluated and scored by a board-certified veterinary pathologist (J.M.S), who was blinded to genotype and experimental manipulation for all groups except for the initial group of mice subjected to phenotyping of multiple organs including decalcified cross section of skull, lungs, heart, kidney, liver, spleen, pancreas, lymph nodes, reproductive tract, stomach, and intestines. These juvenile mice evaluated initially in an unblinded fashion were re-scored blindly prior to manuscript preparation. For subsequent mice, histological analysis was focused on the kidney, liver, and spleen.

For the kidney, expansion of the glomerular mesangial matrix was scored from 0-4, with 0 = normal; 1 = minimal; 2 = mild; 3 = moderate (with accompanying tubular protein); and 4 = severe histological changes. An extent score was also given for the kidney, with 1 representing <10%; 2 = 10-32%; 3 = 33-65%; and 4 representing > 66% of glomeruli affected. For the liver, microvesicular and lesser macrovesicular cytoplasmic vacuolation were scored from 0-5, with 0 = normal; 1 = minimal changes affecting only a small region (< 5%) of the liver; 2 = mild changes throughout the liver but without enlargement of hepatocytes, coalescing lesions, or necrosis; 3 = mild to moderate cytoplasmic vacuolation throughout liver with enlargement of hepatocytes but no necrosis or loss of parenchyma; 4 = moderate, coalescing throughout liver with multifocal mild regions of loss of parenchyma or necrosis; and 5 = severe with moderate multifocal regions of cavitation and necrosis. Liver inflammation was also scored on a 0-5 scale with 0 = fewer than 2 small microgranulomas per section; 1 = minimal scattered inflammation or microgranulomas; 2 = mild (less than 5% of parenchyma involved); 3 = mild to moderate (11-30%); 4 = moderate (31-60%); and 5 = severe (affecting >60% of parenchyma). Lymphoid depletion of the spleen was scored on a scale of 0-3 with 0 = none; 1 = mild; 2 = moderate; 3 = severe.

Representative images were taken using NIS-Elements BR 3.2 64-bit and plated in Adobe Photoshop Elements. Image white balance, lighting, brightness and contrast were adjusted using auto corrections applied to the entire image. Final magnification is stated.

For measurement of alkaline phosphatase (ALP) and albumin in serum, blood was collected by cardiac puncture from mice at 23 days of age and stored in SST Tubes at 4C until analysis. Samples were run at 2x dilution on a Siemens Atellica 1600 (Siemens Healthineers, Germany).

### mRNASeq and Analysis

Total RNA was added directly to lysis buffer from the SMART-Seq v4 Ultra Low Input RNA Kit for Sequencing (Takara), and reverse transcription was performed followed by PCR amplification to generate full-length amplified cDNA. Sequencing libraries were constructed using the NexteraXT DNA sample preparation kit (Illumina) to generate Illumina-compatible barcoded libraries. Libraries were pooled and quantified using a Qubit^®^ Fluorometer (Life Technologies). Dual-index, single-read sequencing of pooled libraries was carried out on a HiSeq2500 sequencer (Illumina) with 58-base reads, using HiSeq v4 Cluster and SBS kits (Illumina) with a target depth of 5 million reads per sample.

Reads were processed using workflows managed on the Galaxy platform. Reads were trimmed by 1 base at the 3’ end, and then trimmed from both ends until base calls had a minimum quality score of at least 30 (Galaxy FASTQ Trimmer tool v1.0.0). FastqMcf (v1.1.2) was used to remove any remaining adapter sequence. To align the trimmed reads, we used the STAR aligner (v2.4.2a) with the GRCm38 reference genome and gene annotations from ensembl release 91. Gene counts were generated using HTSeq-count (v0.4.1). Quality metrics were compiled from PICARD (v1.134), FASTQC (v0.11.3), Samtools (v1.2), and HTSeq-count (v0.4.1). Libraries with less than 2 million mapped reads per sample were not included in the analysis, and analysis was restricted to genes with non-zero counts in all libraries (*n*=8,843 genes). We carried out normalization and tested for differential expression using DESeq2 (Love et al., 2014). Gene set enrichment analysis was carried out using fgsea (Koretkevich et al., 2019), ranking genes by the sorted p values from the differential expression test for each tissue. Gene sets tested included all GO categories, Kegg pathway gene sets, Hallmark gene sets, and Reactome gene sets. We also included the manually curated ISR target gene set as defined previously (Wong et al., 2019). In total, 7084 gene sets were evaluated, and multiple testing correction was done using the Benjamini-Hochberg method.

### Quantitative RT-PCR

Mice were euthanized by CO_2_ asphyxiation and cardiac puncture. Organs were immediately immersed in Trizol and frozen at −80C until later processing. To extract RNA, organs were mashed on ice, then resuspended in Trizol by passage through 25G needles. Samples were spun down briefly (5 min at 5000 g), and the supernatant processed by the Direct-zol RNA MiniPrep kit (Genesee Scientific) per the manufacturer’s instructions with an additional dry spin after disposing of the final wash to prevent EtOH carryover. Complementary DNA (cDNA) was generated using EcoDry double primed premix (Clontech). Transcript expression was measured using the following ThermoFisher TaqMan Gene expression assays: *Ifi27* (Mm00835449_g1), *Irf7* (Mm00516788_m1), *Oas1a* (Mm00836412_m1), *Asns* (Mm00803785_m1), *Cdkn1a* (Mm04205640_g1), and *Hmox1* (Mm00516005_m1). Samples were assayed in duplicate on a real-time qPCR cycler (CFX96, BioRad).

### *In vitro* treatments

Bone marrow was harvested from age-matched mice and grown/differentiated in RPMI supplemented with 10% fetal calf serum, L-glutamine, penicillin/streptomycin, sodium pyruvate, HEPES, and M-CSF for 7 days. 10^5^ BMDM were plated in 12 well plates and rested overnight, then stimulated with 100U/mL recombinant murine IFNβ (Sigma, I9032) or equivalent volume of water for 24 hours. Cells were then harvested in Trizol before RNA purification via Directzol kits, and cDNA generation using Ecodry kits as described above. qPCR was performed using iTaq supermix on the Bio-Rad CFX96 Real-Time System, and primers for IFNB (*Ifnb* Fwd 5’-GCACTGGGTGGAATGAGACTATTG-3’, *IfnB* Rev 5’-TTCTGAGGCATCAACTGACAGGTC-3’) and HPRT (*Hprt* Fwd, 5’-GTT GGATACAGGCCAGACTTTGTTG-3’, *Hprt* Rev, 5’-GAGGGTAGGCTGGCCTATAGGCT-3’.

### 2BAct treatments

*Adar P195A/P195A* mice were intercrossed with *Adar p150+/-* mice. 48 hours after mating, the breeders were placed on 2BAct chow, formulated to achieve a 2Bact concentration of 300 ppm (300 μg 2BAct/g of meal) as previously described (Wong et al., 2019). 2BAct chow was contract manufactured with Envigo. The placebo diet was Teklad 2014 without added compound. Breeders and pups were maintained on 2BAct until termination of survival experiments at 125 days. Mice were weighed at 23 days of age. Littermate mice were harvested for histological and mRNA expression analysis as described above.

### Statistical analysis

Quantitative data were visualized and analyzed using GraphPad Prism software. Differences in survival were assessed by the Log-rank (Mantel-Cox) Test. Observed and expected birth rates by genotypes were compared using the Chi-square test. Weights and mRNA expression measured by qPCR were compared between genotypes within each cross using unpaired t-tests. Significance is indicated as follows in all figures: ns=not significant, *p<0.05, **p<0.01, ***p<0.001, ****p<0.0001.

## Acknowledgements

We thank Stuart Orkin, Michael Oldstone, Michael Gale, Jr, Gökhan Hotamişligil, and Robert Silverman for providing mice used in this study. We are grateful to Leonel Joannas for expertise in the generation of the *Adar P195A* mice, to Sarah Miller for expert mouse colony management, and to all members of the Stetson lab for insightful discussions. This work was supported by NIH R01 AI084914 (D.B.S.); and T32 AI106677 and NIH F31 AI40432 (M.M.) D.B.S. is a Faculty Scholar of the Howard Hughes Medical Institute.

## Author Contributions

M.M. and D.B.S. conceived the study. M.M. performed the experiments. J.M.S. performed the histological analyses. C.C. performed the mRNA-Seq analyses. J.H.-M. generated the *Adar P195A* knockin mice. C.S. coordinated the 2BAct experiments and mRNA-Seq analyses. M.M. and D.B.S. wrote the manuscript.

---

**Supplemental Figure 1.**
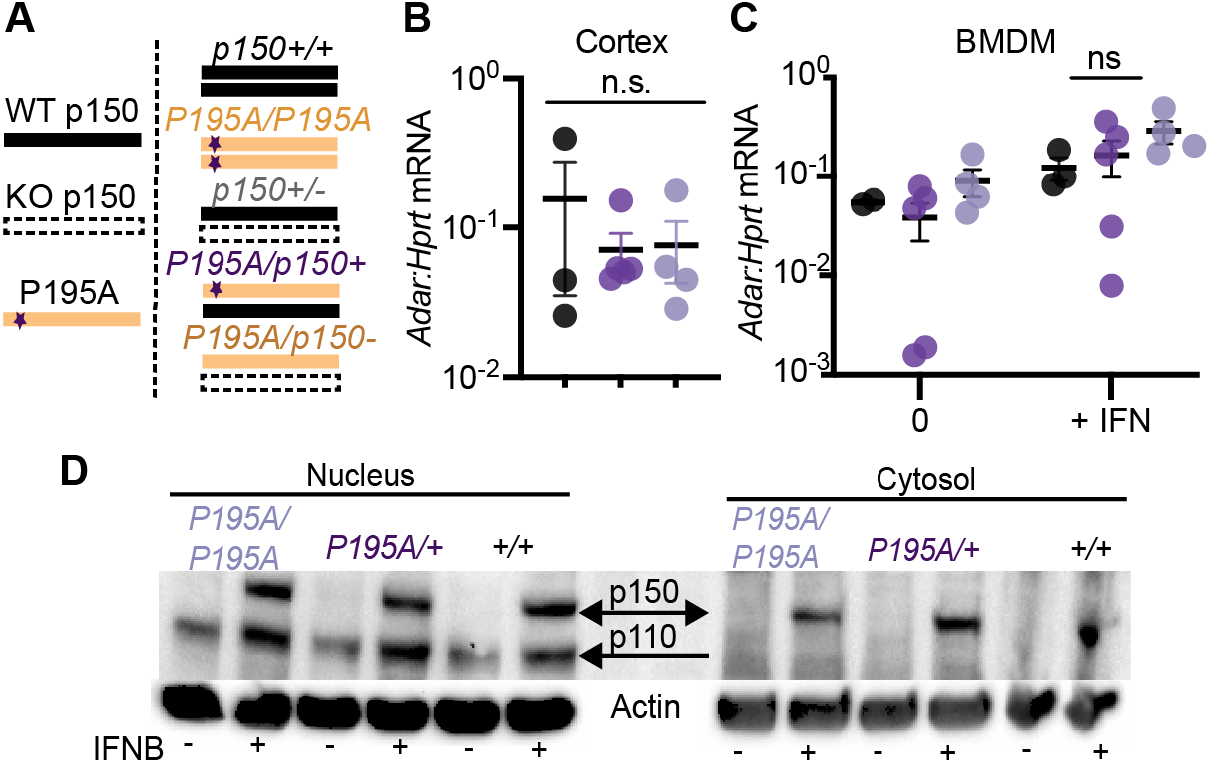
Characterization of the *Adar P195A* knockin mouse model. (A) Schematics representing the different *Adar p150* alleles, and different *Adar P195A/p150* genotypes. (B) Expression of *Adar* mRNAs in the cortex of *Adar P195A/P195A, Adar P195A/+* and *Adar+/+* mice. (C) Expression of *Adar* mRNAs in BMM from *Adar P195A/P195A, Adar P195A/+* and *Adar+/+* mice, with and without 24 hour treatment with recombinant IFNβ. Bars represent mean and SEM. (D) Western blot for ADAR1 in the cytosol and nucleus of primary MEFs of the indicated genotypes with and without 24 hour treatment with recombinant IFNβ.

**Supplemental Figure 2.**
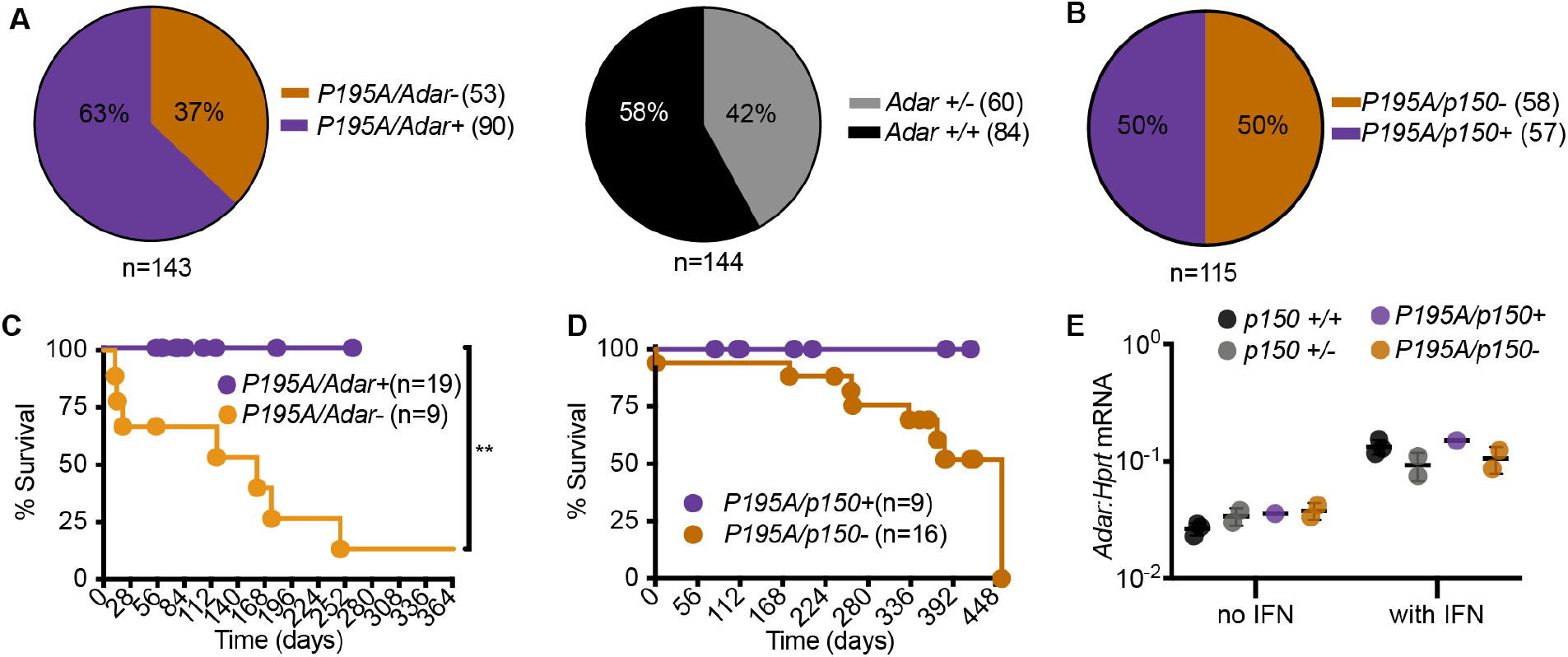
Features of *Adar P195A* mice crossed to *Adar* null alleles. (A) Percentage of mice of the indicated genotype born from crosses of *Adar P195A/P195A* and *Adar+/-* mice (left panel), and *Adar+/+* and *Adar+/-* mice (right panel). Number of each genotype is indicated in parentheses. *Adar+/-* mice are born at a lower than Mendelian frequency, as has been previously observed (Pestal et al., 2015). (B) Percentage of mice of the indicated genotype born from crosses of *Adar P195A/P195A* and *Adar p150+/-* mice (right panel). (C) Survival of mice of the indicated *Adar* genotype on an *Ifih1+/-* background, revealing partial rescue from mortality. (D) Survival of mice of the indicated *Adar p150* genotype on an *Ifih1+/-* background, revealing partial rescue from mortality. (E) Expression of *Adar* mRNA transcript in independently derived primary MEFs of the indicated genotypes, with and without 24 hours of treatment with recombinant mouse IFNβ. Bars represent mean and SEM.

**Supplemental Figure 3.**
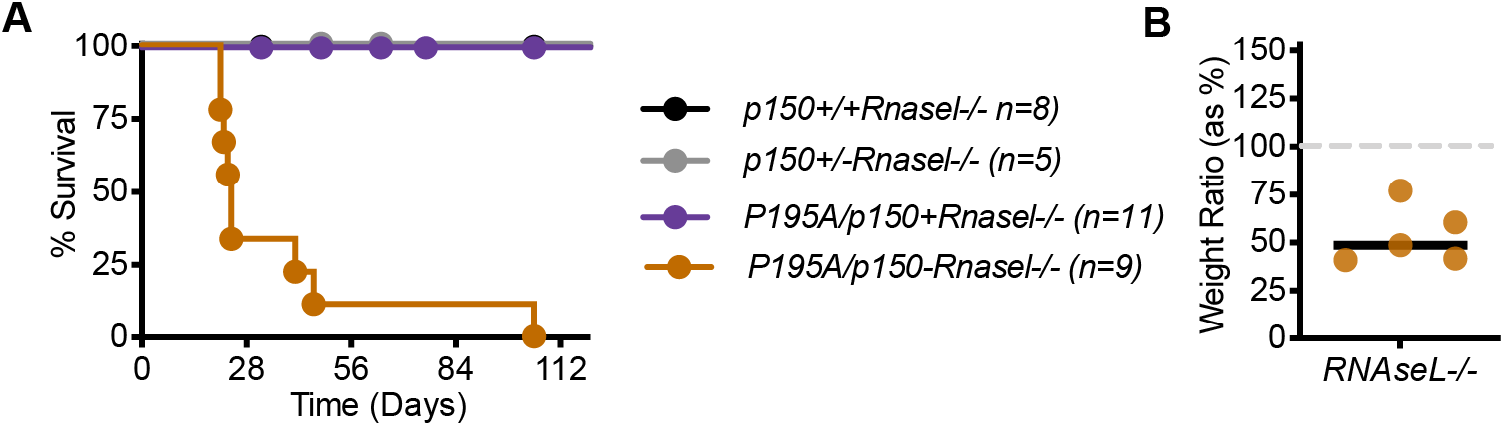
RNase L deficiency does not rescue Adar P195A/p150- mice. (A) Survival of *Rnasel-/-* mice of the indicated genotype: *Adar p150+/+* (n=8), Adar *p150+/-* (n=5), *Adar P195A/p150+* (n=11), *Adar P195A/p150*- (n=9). (B) Weights, measured at 23 days, of *Adar P195A/p150-Rnasel-/-* mice, as a percentage of the average weight of age- and sex-matched *Adar P195A/p150+Rnasel-/-* control mice.

